# Ribosome profiling reveals post-translational signaling mechanisms drive the retrograde enhancement of presynaptic efficacy

**DOI:** 10.1101/158303

**Authors:** Xun Chen, Dion K. Dickman

## Abstract

Presynaptic efficacy can be modulated by retrograde control mechanisms, but the nature of these complex signaling systems remain obscure. We have developed and optimized a tissue specific ribosome profiling approach in *Drosophila.* We first demonstrate the ability of this technology to define genome-wide translational regulations. We then leverage this technology to test the relative contributions of transcriptional, translational, and post-translational mechanisms in the postsynaptic muscle that orchestrate the retrograde control of presynaptic function. Surprisingly, we find no changes in transcription or translation are necessary to enable retrograde homeostatic signaling. Rather, post-translational mechanisms appear to ultimately gate instructive retrograde communication. Finally, we find that a global increase in translation induces adaptive responses in both transcription and translation of protein chaperones and degradation factors to promote cellular proteostasis. Together, this demonstrates the power of ribosome profiling to define transcriptional, translational, and post-translational mechanisms driving retrograde signaling during adaptive plasticity.

**AUTHOR SUMMARY:** Recent advances in next-generation sequencing approaches have revolutionized our understanding of transcriptional expression in diverse systems. However, transcriptional expression alone does not necessarily report gene translation, the process of ultimate importance in understanding cellular function. To circumvent this limitation, biochemical tagging of ribosomes and isolation of ribosomally-associated mRNA has been developed. However, this approach, called TRAP, has been shown to lack quantitative resolution compared to a superior technology, ribosome profiling, which quantifies the number of ribosomes associated with each mRNA. Ribosome profiling typically requires large quantities of starting material, limiting progress in developing tissue-specific approaches. Here, we have developed the first tissue specific ribosome profiling system in *Drosophila* to reveal genome-wide changes in translation. We first demonstrate successful ribosome profiling from a specific tissue, muscle, with superior resolution compared to TRAP. We then use transcriptional and ribosome profiling to define transcriptional and translational adaptions necessary for synaptic signaling at the neuromuscular junction. Finally, we utilize ribosome profiling to demonstrate adaptive changes in cellular translation following cellular stress to muscle tissue. Together, this now enables the power of *Drosophila* genetics to be leveraged with translational profiling in specific tissues.

## INTRODUCTION

Synapses have the capacity to adaptively modulate the efficacy of neurotransmission to maintain stable levels of functionality, counteracting perturbations that would otherwise impair communication in the nervous system. These robust homeostatic mechanisms stabilize synaptic strength in both the central and peripheral nervous systems, and have been demonstrated to exist in invertebrates to humans (Davis and Muller, 2015; Pozo and Goda, 2010; Turrigiano, 2012). In each system, disruption of synaptic transmission leads to compensatory changes in postsynaptic receptor trafficking or presynaptic efficacy that restores baseline levels of activity (Davis, 2013; Turrigiano, 2008). The importance of homeostatic signaling is underscored by links with a variety of neurological and neuropsychiatric diseases, including epilepsy, schizophrenia, and autism (Meier et al., 2014; Nelson and Valakh, 2015; Wondolowski and Dickman, 2013). Although a perturbation to synaptic activity is clearly required for the induction of homeostatic synaptic plasticity, the mechanisms that respond to this perturbation to induce intrinsic and trans-synaptic homeostatic signaling remain largely unknown.

The *Drosophila* neuromuscular junction (NMJ) has been established as a powerful system to reveal the genes and mechanisms involved in the homeostatic control of synaptic strength (Frank, 2014; Frank et al., 2006; Petersen et al., 1997). At this glutamatergic synapse, genetic or pharmacological perturbations that disrupt postsynaptic neurotransmitter receptors induce a retrograde signaling system that ultimately potentiates presynaptic release, restoring baseline levels of synaptic transmission (Frank et al., 2006; Haghighi et al., 2003; Petersen et al., 1997). Specifically, reduction in the amplitude of miniature excitatory postsynaptic potentials (mEPSPs) are observed in response to loss of the GluRIIA subunit (Fig. 1A). However, excitatory postsynaptic receptor amplitude (EPSP) is maintained at wild-type levels due to an enhancement in the number of synaptic vesicles released (quantal content). This process is referred to as presynaptic homeostatic potentiation (PHP) because the expression mechanism of this form of plasticity is a presynaptic increase in neurotransmitter release. Recent forward genetic screening and candidate approaches have revealed the identity of several genes and effector mechanisms in the presynaptic neuron required for the homeostatic potentiation of presynaptic release (Dickman and Davis, 2009; Dickman et al., 2012; Frank et al., 2009; Genc et al., 2017; Harris et al., 2015; Kiragasi et al., 2017; Muller and Davis, 2012; Muller et al., 2015; Muller et al., 2012; Muller et al., 2011; Tsurudome et al., 2010; Wang et al., 2016; Younger et al., 2013). However, in contrast to our understanding of the genes and mechanisms involved in PHP expression in the presynaptic neuron, very little is known about the postsynaptic signaling system that transduces a reduction in glutamate receptor function into a retrograde signal that instructs an adaptive increase in presynaptic release.

**Fig 1:**
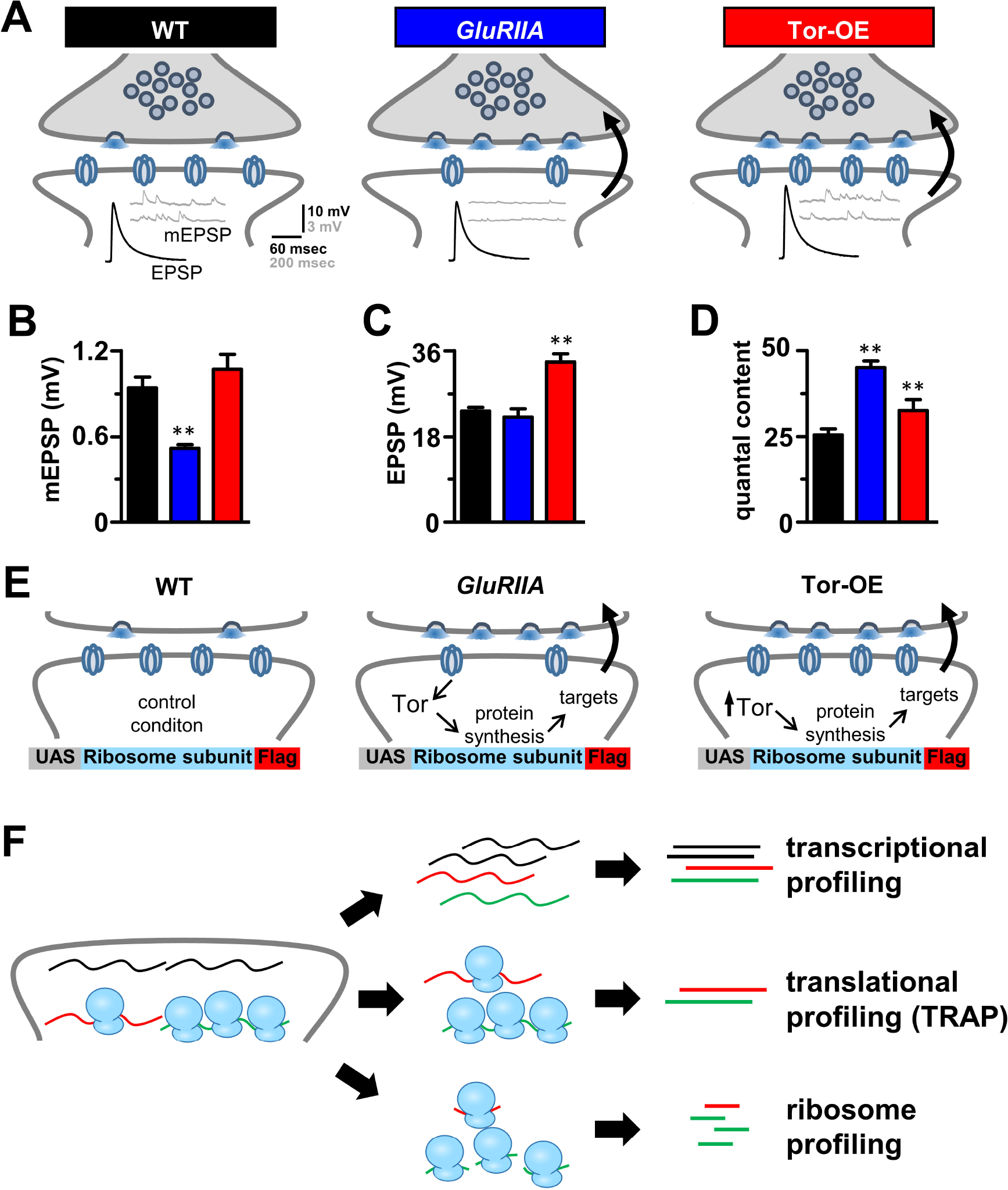
Schematic detailing transcriptional and translational profiling of retrograde signaling at the *Drosophila* NMJ. **(A)** Schematic illustrating synaptic transmission at the *Drosophila* NMJ. Representative EPSP and mEPSP electrophysiological traces in wild type (*w*^*1118*^; n=6), *GluRIIA* mutants (*w*;*GluRIIA*^*sp16*^; n=6), and overexpression of *Tor* in the postsynaptic muscle (Tor-OE: *w*;*UAS-Tor-myc/+;BG57-Gal4/+;* n=6). Note that while mEPSP amplitudes are reduced in *GluRIIA* mutants, EPSP amplitudes remain the same as wild type because of a homeostatic increase in presynaptic release (quantal content). Tor-OE does not change mEPSP amplitude, but retrograde homeostatic signaling is induced, leading to increased EPSP amplitude and quantal content. Quantification of mEPSP amplitude **(B)**, EPSP amplitude **(C)**, and quantal content **(D)** for the indicated genotypes. **(E)** Schematic illustrating the putative role of protein synthesis in retrograde homeostatic signaling and the design of ribosome tagging to isolate postsynaptic RNA. **(F)** Schematic representing the workflow for transcriptional profiling, translational profiling using TRAP (translating ribosome affinity purification), and ribosome profiling. Student’s t test was used to compare *GluRIIA* and Tor-OE to wild type; **=p<0.01.

Modulation of protein synthesis through the Target of Rapamycin (Tor) pathway was recently implicated in the postsynaptic signaling system controlling PHP. Genetic loss of the postsynaptic glutamate receptor subunit *GluRIIA* at the *Drosophila* NMJ leads to a chronic reduction in postsynaptic excitability, but normal synaptic strength due to a homeostatic increase in presynaptic release (Petersen et al., 1997). However, pharmacologic or genetic inhibition of postsynaptic protein synthesis through the Tor pathway and associated translational modulators disrupts the expression of PHP (Kauwe et al., 2016; Penney et al., 2012; Penney et al., 2016). Further, a constitutive increase in muscle protein synthesis through postsynaptic overexpression of Tor was sufficient to trigger the retrograde enhancement of presynaptic release without any perturbation to glutamate receptors (Penney et al., 2012; Penney et al., 2016). While these results establish some of the first insights into the postsynaptic signal transduction system controlling retrograde PHP signaling, the putative translational targets involved, and to what extent transcriptional and/or post-translational mechanisms contribute to PHP signaling, remain unknown.

Recent advances in next-generation sequencing have enabled the ability to quantify genome-wide changes in RNA expression, without pre-existing knowledge, at unprecedented resolution (Chen et al., 2016; Ozsolak and Milos, 2011; Wang et al., 2009). In addition, biochemical tagging of ribosomes has emerged as a powerful way to separate and define the actively translating mRNA pool from overall mRNA expression (Chen and Rosbash, 2017; Heiman et al., 2008; Huang et al., 2013; Sanz et al., 2009; Thomas et al., 2012; Yang et al., 2005; Zhang et al., 2016), a technique termed TRAP (Translating Ribosome Affinity Purification) followed by RNA-seq. Although this approach provides important insights into translational regulation, it lacks the resolution to differentiate between mRNA populations associated with few or high numbers of ribosomes, a distinction that can have major consequences for accurately defining translational rates (Chekulaeva and Landthaler, 2016; Heiman et al., 2014). This limitation was recently overcome through the development of a technique called “ribosome profiling”, which quantifies only mRNA fragments that are protected by ribosomes (“ribosome footprints”). This enables the quantitative analysis of the number of ribosomes associated with each mRNA transcript, and is even capable of defining regions within RNA transcripts of ribosome association (Ingolia, 2016; Ingolia et al., 2009; Li et al., 2014). This technology has been used to reveal genome-wide adaptations to translation that would not have been observed from transcriptional or translational profiling (TRAP) approaches alone (Cho et al., 2015; Dunn et al., 2013; Gonzalez et al., 2014; Ingolia et al., 2009; Ingolia et al., 2011; Jeong et al., 2016). However, despite the potential of ribosome profiling, this approach has not been developed for tissue-specific applications in *Drosophila,* nor brought to the study of retrograde homeostatic signaling.

We have developed and optimized a streamlined system for ribosome profiling of specific tissues in *Drosophila.* We first demonstrate the success of this approach in defining translational regulation in the larval muscle, and reveal dynamics in translation that are distinct from overall transcriptional expression. Next, we highlight the superior sensitivity of ribosome profiling in reporting translational regulation over the conventional TRAP method. We go on to use ribosome profiling to assess the contributions of transcriptional, translational, and post translational mechanisms in the postsynaptic muscle that drive the retrograde signaling system underlying presynaptic homeostatic potentiation. Unexpectedly, we find no changes in postsynaptic transcription or translation following PHP signaling. Instead, post-translational mechanisms appear to be necessary, which can transform an overall increase in cellular translation into a specific, instructive retrograde signal. Finally, our analysis also revealed adaptive changes in both transcription and translation in the postsynaptic cell in response to chronically elevated Tor-mediated translation, including increased expression of protein chaperones, ubiquitin ligases, and ribosome biogenesis factors. Thus, we have established a tissue-specific ribosome profiling approach in *Drosophila* and used it to define transcriptional, translational, and post-translational contributions to retrograde signaling during synaptic plasticity.

## RESULTS

### A strategy to profile postsynaptic transcription and translation adaptations that drive retrograde PHP signaling

To assess the postsynaptic retrograde signaling systems that drive presynaptic homeostatic potentiation (PHP) at the *Drosophila* NMJ, we focused on three genetic conditions (Fig. 1A). First is the wild-type control genotype (*w^1118^*), which serves as the control condition in which PHP is not induced or expressed. Second, null mutations in the postsynaptic glutamate receptor subunit *GluRIIA* (*GluRIIA^SP16^*) lead to a chronic reduction in mEPSP amplitude (Petersen et al., 1997). However, EPSP amplitudes are maintained at wild-type levels due to a homeostatic increase in presynaptic release (quantal content) following retrograde signaling from the muscle (Fig. 1A-D). This serves as one condition in which we hypothesized that gene transcription, translation, and/or post-translational changes may have occurred in response to loss of *GluRIIA,* triggering the induction of retrograde signaling that drives PHP. Indeed, in *GluRIIA* mutants, genetic disruption of the translational regulator *Target of rapamycin* (*Tor*) blocks PHP expression, resulting in no change in quantal content and a concomitant reduction in EPSP amplitude (Penney et al., 2012). Finally, postsynaptic overexpression of *Tor* in an otherwise wild-type muscle (Tor-OE: *UAS-Tor-myc/+;BG57-Gal4/+*) is sufficient to trigger PHP signaling, leading to increased presynaptic release and EPSP amplitude with no change in mEPSP or glutamate receptors (Fig. 1A-D; (Penney et al., 2012)). Tor-OE therefore served as the final genotype in which PHP signaling was induced through *Tor* overexpression without any perturbation of postsynaptic glutamate receptors. We hypothesized that shared changes in translation, and perhaps even transcription, between *GluRIIA* mutants and Tor-OE would illuminate the nature of the postsynaptic transduction system underlying homeostatic retrograde signaling at the *Drosophila* NMJ.

To define genome-wide changes in mRNA transcription and translation in the postsynaptic muscle that may be necessary for PHP signaling, we sought to purify RNA from 3^rd^instar larvae muscle in wild type, *GluRIIA* mutants, and Tor-OE (Fig. 1E). We then sought to define mRNA expression through three methods: Transcriptional profiling, translational profiling using translating ribosome affinity purification (TRAP), and ribosome profiling (Fig. 1F). First, transcriptional profiling of total mRNA expression can be performed by isolating RNA from dissected third instar muscle and prepared for RNA-seq through standard methods (Brown et al., 2014; Mortazavi et al., 2008). To define translational changes, we engineered an affinity tag on a ribosome subunit under control of the upstream activating sequence (UAS), which enables tissue-specific expression (Fig. 1E). This biochemically tagged ribosome could therefore be expressed in muscle to purify ribosomes, then processed to sequence only mRNA sequences associated with or protected by ribosomes (Fig. 1F). Affinity tagging of ribosomes enabled us to perform translational profiling (TRAP: Translating Ribosome Affinity Purification), an established technique capable of detecting ribosome-associated mRNA (Chen and Rosbash, 2017; Heiman et al., 2014; Heiman et al., 2008; Huang et al., 2013; Zhang et al., 2016). Finally, we reasoned that this approach could be optimized to enable ribosome profiling, which has been used successfully to determine changes in translational rates, with superior sensitivity over TRAP, in a variety of systems (Dunn et al., 2013; Ingolia et al., 2009; Ingolia et al., 2011; Jeong et al., 2016). However, ribosome profiling has not been developed for use in specific *Drosophila* tissues. Our next objective was to optimize a tissue-specific ribosome profiling approach.

### Optimization of a tissue-specific ribosome profiling approach in *Drosophila*

Ribosome profiling is a powerful approach for measuring genome-wide changes in mRNA translation rates. However, high quantities of starting material is necessary to obtain sufficient amounts of ribosome protected mRNA fragments for the subsequent processing steps involved (Brar and Weissman, 2015). Since this approach has not been developed for *Drosophila* tissues, we first engineered and optimized the processing steps necessary to enable highly efficient affinity purification of ribosomes and ribosome protected mRNA fragments by incorporating ribosome affinity purification into the ribosome profiling protocol.

Although tissue-specific ribosome affinity purification strategies have been developed before in *Drosophila* (Thomas et al., 2012; Zhang et al., 2016), these strategies have not been optimized to meet the unique demand necessary for ribosome profiling. We thus set out to develop and optimize a new ribosome affinity purification strategy that enables the efficient purification and processing of ribosomally-protected mRNA. First, we generated transgenic animals that express a core ribosome subunit in frame with a biochemical tag (*3×Flag*) under *UAS* control to enable expression of this transgene in specific *Drosophila* tissues (Fig. 1E and 2A). Although previous approaches in *Drosophila* have targeted the same ribosome subunit (RpL10a) with different tags (Huang et al., 2013; Thomas et al., 2012; Zhang et al., 2016), we found these to be sub-optimal for the efficiency necessary for ribosome profiling (data not shown). Therefore, based on high resolution crystal structures of eukaryotic ribosomes (Ben-Shem et al., 2011; Khatter et al., 2015), we selected an alternative ribosomal protein from the large and small subunits expected to have C terminals exposed on the ribosome surface. We cloned the *Drosophila* homologs of these subunits, *RpL3* and *RpS13,* in frame with a C-terminal *3×Flag* tag and inserted this sequence into the pACU2 vector for high expression under *UAS* control (Han et al., 2011). We then determined whether intact ribosomes could be isolated in muscle tissue following expression of the tagged ribosomal subunit. We drove expression of *UAS-RpL3-Flag* or *UAS-RpS13-Flag* with a muscle-specific *Gal4* driver (*BG57-Gal4*) and performed anti-Flag immunoprecipitations (Fig. 2A). An array of specific bands were detected in a Commassie stained gel from the RpL3-Flag and RpS13-Flag immunoprecipitations, but no such bands were observed in lysates from wild type (Fig. 2B). Importantly, identical sized bands were observed in immunoprecipitates from both RpL3-Flag and RpS13-Flag animals, matching the expected distribution of ribosomal proteins (Anger et al., 2013). The RPL3-Flag immunoprecipitation showed the same distribution as RpS13 but higher band intensity, indicating higher purification efficiency, so we used *RpL3-Flag* transgenic animals for the remaining experiments. In addition to ribosomal proteins, the other major constituent of intact ribosomes is ribosomal RNA. Significant amounts of ribosomal RNA were detected in an agarose gel from RpL3-Flag immunoprecipitates (Fig. 2C), providing additional independent evidence that this affinity purification strategy was efficient at purifying intact ribosomes.

**Fig 2:**
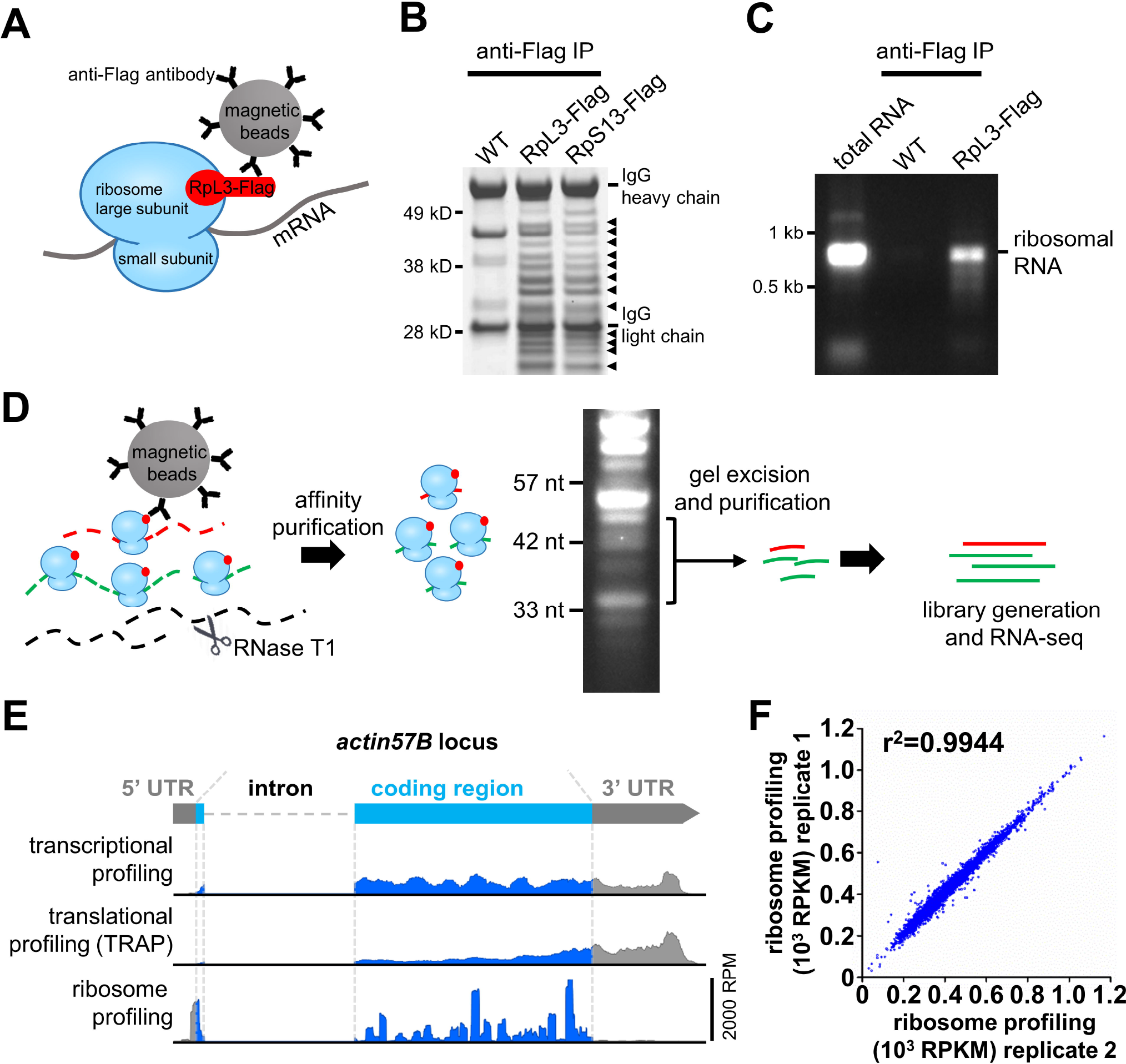
Development and verification of an optimized ribosome profiling protocol in *Drosophila.* **(A)** Schematic illustrating the ribosome affinity purification strategy. A tagged ribosome subunit (RpL3-Flag) is expressed and incorporated into ribosomes. Magnetic beads coated with anti-flag antibodies are used to immunoprecipitate ribosomes along with associated mRNA. **(B)** Anti-flag immunoprecipitation from wild-type control, postsynaptic expression of RpL3-Flag (*w*;*BG57-Gal4/UAS-RpL3-3xflag*), and postsynaptic expression of RpS13-Flag (*w*;*BG57-Gal4/UAS-RpS13-3xflag*) in third-instar larval muscle. Sample was run on an SDS-PAGE gel and Commassie stained. The expected distribution of ribosomal proteins are present in RpL3-Flag and RpS13-Flag samples (noted by arrowheads), but not observed in wild-type controls. **(C)** Total RNA was extracted from anti-flag immunoprecipitations from wild type and RpL3-Flag larval muscle tissue and run on an agarose gel. Ribosomal RNA is present in RpL3-Flag RNA samples but absent in wild-type samples. Total RNA extracted from wild type whole larvae was loaded to show the position of ribosomal RNA. **(D)** Workflow for ribosome profiling strategy. **(E)** Representative RNA-seq mapping of the *actin57B* locus from transcriptional, translational (TRAP), and ribosome profiling. Note that ribosome profiling reads predominantly map to 5’UTR and coding regions, and are absent from the 3’UTR. RPM: reads per million total mapped reads. **(F)** Replicate ribosome profiling sequencing demonstrates highly reproducible results.

Next, we tested the ability of RpL3-Flag to functionally integrate into intact ribosomes. We generated an *RpL3-Flag* transgene under control of the endogenous promotor (*genomic-RpL3-Flag;* Fig. S1A). This transgene was able to rescue the lethality of homozygous *RpL3* mutations (Fig. S1A), demonstrating that this tagged ribosomal subunit can integrate and function in intact endogenous ribosomes, effectively replacing the endogenous untagged RpL3 protein. Further, anti-Flag immunostaining of *UAS-RpL3-Flag* expressed in larval muscle showed a pattern consistent with expected ribosome distribution and localization (Fig. S1B). Thus, biochemical tagging of RpL3 does not disrupt its localization or ability to functionally integrate into endogenous ribosomes.

Finally, we developed and optimized a method to process the isolated ribosomes to generate only ribosome protected mRNA fragments to be used for RNA-seq analysis. First, we digested the tissue lysate with RNaseT1, an enzyme that cuts single stranded RNA at G residues (Fig. 2D). Following digestion, we ran RNA on a high percentage PAGE gel, excising the mRNA fragments protected from digestion by ribosome binding (30-45 nucleotides in length; Fig. 2D). Sequencing of this pool of RNA demonstrated that the vast majority of reads mapped to the 5’UTR and coding regions of mRNA transcripts, with very few reads mapping to the 3’UTR of mRNA transcripts (Fig. 2E), where ribosomes are not expected to be associated. This coverage map also revealed heterogeneous distributions on mRNA transcripts with irregular and prominent peaks, as expected, which are indicative of ribosome pause sites on mRNA (Fig. 2E; (Li et al., 2012)). In contrast, RNA-seq reads for transcriptional and translational profiling using TRAP mapped to the entire mRNA transcript with relatively even coverage (Fig. 2E). Importantly, replicate experiments demonstrated that this protocol generated highly reproducible measures of relative protein synthesis rates, defined by mRNA ribosome density, or the number of ribosome profiling Reads Per Kilobase of exon per Million mapped reads (RPKM, also referred to as ribosome profiling expression value; Fig. 2F). Thus, expression of *RpL3-Flag* enables the purification of ribosomes from specific tissues in *Drosophila,* and further processing reproducibly generates ribosome protected mRNA fragments, which correlate with protein synthesis rates (Li et al., 2014).

### Ribosome profiling is more sensitive in detecting translational regulation compared to translational profiling (TRAP)

Translation can differ in significant ways from overall transcriptional expression through modulations in the degree of ribosome association with each mRNA transcript, in turn suppressing or enhancing protein synthesis rates (Chekulaeva and Landthaler, 2016; Kong and Lasko, 2012). Translation efficiency is a measure of these differences, defined as the ratio of translational to transcriptional expression (Ingolia et al., 2009). Hence, translation efficiency (TE) reflects the enhancement or suppression of translation relative to transcriptional expression due to various translational control mechanisms (Jackson et al., 2010; Kong and Lasko, 2012). Although both translational (TRAP) and ribosome profiling approaches can report TE, ribosome profiling should, in principle, exhibit superior sensitivity in revealing translational dynamics. We therefore compared translational and ribosome profiling directly to test this prediction.

We compared translation efficiency by comparing TRAP and ribosome profiling to transcriptional profiling in wild-type muscle. In particular, we tested whether differences were apparent in the number of genes revealed to be translationally suppressed or enhanced relative to transcriptional level through ribosome profiling compared to TRAP. We first analyzed the extent to which ribosome profiling and TRAP measurements correlate with transcriptional profiling by plotting the ribosome profiling and TRAP expression values as a function of transcriptional profiling (Fig. 3A,B; see materials and methods). A low correlation would indicate more translational regulation is detected, while a high correlation is indicative of less translational regulation. This analysis revealed a low correlation between ribosome profiling and transcriptional profiling (correlation of determination *r*^2^=0.100; Fig. 3A), while a relatively high correlation was observed between TRAP and transcriptional profiling (*r*^2^=0.617; Fig. 3B). Further, we subdivided all measured genes into three categories: high TE, medium TE, and low TE. These groups were based on translation efficiency as measured by ribosome profiling or TRAP, with high TE genes having a TE value >2, representing genes that are translationally enhanced relative to transcriptional level; low TE genes having a TE value <0.5, representing genes translationally suppressed relative to transcriptional level; and medium TE genes having a TE between 0.5 and 2; representing genes not under strong translational regulation. This division revealed a higher number of genes in the high and low TE groups detected by ribosome profiling compared to TRAP (Fig. 3C). Together, these results are consistent with ribosome profiling detecting more genes under translational regulation compared to TRAP.

**Fig 3:**
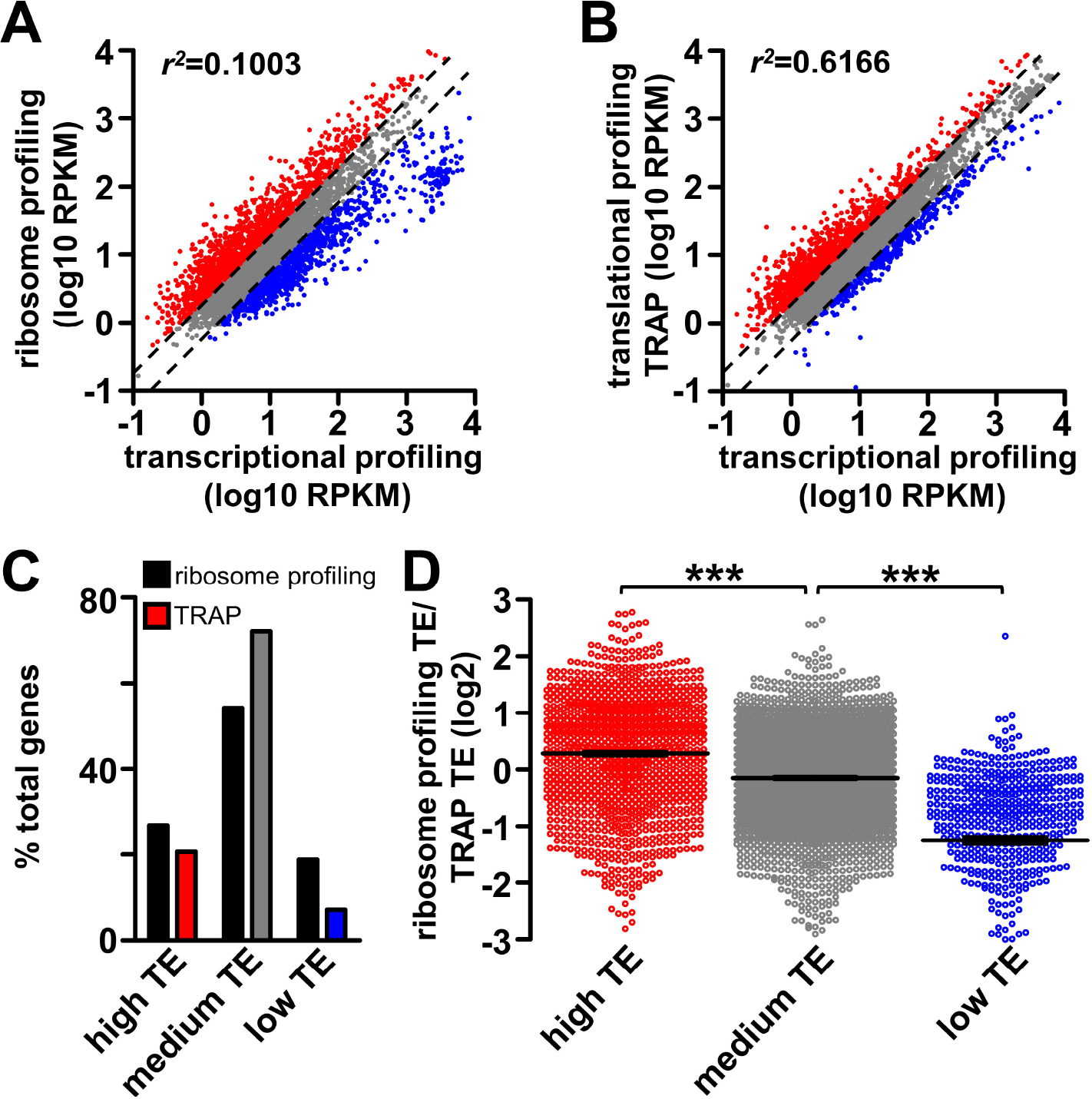
Comparison of translational and ribosome profiling from *Drosophila* larval muscle. **(A)** Plot of ribosome profiling RPKM as a function of transcriptional profiling RPKM for all muscle genes in wild type. Genes with high translation efficiency (TE; TE>2) or low TE (TE<0.5) are labeled in red and blue respectively. Genes with medium TE (TE between 0.5 and 2, indicated by the two dash lines) are labeled in grey. **(B)** Plot of translational profiling (TRAP) RPKM as a function of transcriptional profiling RPKM for all muscle genes in wild type. The same color coding scheme is used as in (A). **(C)** Graph showing percentage of total muscle genes that are in the high TE, medium TE or low TE group based on ribosome profiling or TRAP. Note that a lower percentage of genes are revealed to have high or low TE with TRAP compared to ribosome profiling. **(D)** Plot of translational efficiency of genes defined by ribosome profiling as a ratio of TRAP in three categories: high TE (ribosome profiling and TRAP TE average>2), medium TE (TE average between 0.5 and 2), and low TE (TE average<0.5). Note that ribosome profiling reveals higher TE for high TE genes, and lower TE for low TE genes compared to TRAP. ***=p<0.001; one-way ANOVA with post hoc Bonferroni’s test.

We next investigated the genes under significant translational regulation (genes with high TE or low TE), detected through either ribosome profiling or TRAP, to determine whether differences exist in the amplitude of translational regulation. Specifically, genes were divided into the three categories mentioned above based on the average translation efficiency measured by ribosome profiling and TRAP. We then determined the TE value ratio from ribosome profiling compared to TRAP within the three categories. A ratio above 0 (log2 transformed) in the high TE group indicates a more sensitive reporting of translation for ribosomal profiling, while a ratio below 0 in the low TE group would also indicate superior sensitivity for the ribosomal profiling approach. This investigation revealed an average ratio of 0.28 within the high TE group, -0.15 within the medium TE group, and -1.25 within the low TE group (Fig. 3D). This analysis demonstrates that ribosome profiling is at least 22% more sensitive in detecting high TE, and 138% more sensitive in detecting low TE in comparison to TRAP. Thus, this characterization demonstrates that ribosome profiling provides a more sensitive and quantitative measurement of translational regulation in comparison to TRAP, validating this approach.

### Transcriptional and ribosome profiling reveals dynamic translational regulation in *Drosophila* muscle

Both subtle and dramatic differences have been observed in rates of mRNA translation relative to transcription, particularly during cellular responses to stress (Dunn et al., 2013; Halbeisen and Gerber, 2009; Spriggs et al., 2010). Having optimized and validated our approach, we went to perform transcriptional and ribosome profiling in *GluRIIA* mutants and Tor-OE in addition to wild type (Table S1). To minimize genetic variation, the three genotypes were bred into an isogenic background, and three replicate experiments were performed for each genotype (see materials and methods). We first determined the total number of genes expressed in *Drosophila* muscle, as assessed through both transcription and ribosome profiling. The fly genome is predicted to encode 15,583 genes (NCBI genome release 5_48). We found 6,835 genes to be expressed in wild-type larval muscle through transcriptional profiling, and a similar number (6,656) through ribosome profiling (Fig. 4A), with ~90% of transcripts being shared between the two lists (Table S2), indicating that the vast majority of transcribed genes are also translated. We found no significant differences in the size of the transcriptome and translatome between wild type, *GluRIIA* mutants, and Tor-OE (Table S2). We then compared the muscle transcriptome to a published transcriptome from the central nervous system (CNS) of third-instar larvae (Brown et al., 2014). This comparison revealed dramatic differences in gene expression between the two tissues (Fig. 4B). In particular, we found several genes known to be enriched in muscle, including *myosin heavy chain, α actinin,* and *zasp52,* to be significantly transcribed and translated in muscle, as expected. In contrast, neural-specific genes such as the active zone scaffold *bruchpilot,* the post-mitotic neuronal transcription factor *elav,* and the microtubule associated protein *tau,* were highly expressed in the CNS but not detected in muscle (Table S3 and data not shown). Together, this demonstrates that the muscle transcriptome and translatome can be defined by the transcriptional and ribosome profiling strategy we developed with high fidelity.

Next, we investigated genome wide translation efficiency distribution in larval muscle, and compared this with gene expression as assessed through transcriptional and ribosome profiling. We first calculated translation efficiency for all genes expressed in larval muscle and compared heat maps of TE to heat maps of the transcription and translation level (Fig. 4C). This revealed a dynamic range of translation efficiency, and a surprising trend of genes with high TE displaying relatively low transcriptional expression levels, while genes with low TE exhibited high transcriptional expression levels (Fig. 4C). We then analyzed the genes categorized as high TE, medium TE and low TE (described above) in more detail, comparing the relative distribution in transcriptional expression. We found this trend to be maintained, in that high TE genes exhibited significantly lower transcriptional expression, while low TE genes were significantly higher in transcriptional expression (Fig. 4D). Together, this implies a general inverse correlation between translational efficiency and transcriptional expression.

Finally, we examined the genes with the most extreme translation efficiency to gain insight into the functional classes of genes that exhibit strong translational control under basal conditions. Interestingly, among the genes with most suppressed translation (100 genes with the lowest translation efficiency), we found a surprisingly high enrichment of genes encoding ribosome subunits and translation elongation factors (Fig. 4E,F; Fig. S2A and Table S3). Indeed, 73 of the 100 genes with the lowest translation efficiency were ribosome subunits, with all subunits exhibiting a consistently low TE, averaging 0.091. In contrast, *RpL3*, the subunit we overexpressed (*UAS-RpL3-Flag*), was a clear outlier compared with the other ribosome subunits, as expected, showing a translation efficiency of 2.85. This overall suppression in TE of ribosome subunits may enable a high dynamic regulatory range, enabling a rapid increase in production of ribosomal proteins under conditions of elevated protein synthesis. Consistent with this idea, we observed a coordinated upregulation of translation efficiency for ribosomal subunits when overall muscle translation is elevated in Tor-OE (Fig. 4H). This is in agreement with previous findings showing ribosome subunits and translation elongation factors as targets for translational regulation by Tor (Jefferies et al., 1994; Thomas et al., 2012). In contrast to the enrichment of ribosome subunits observed in the low TE group, diverse genes were found among the most translationally enhanced group, with genes involved in cellular structure being the most abundant (Fig. 4E,G; Fig. S2B and Table S4). These genes may encode proteins with high cellular demands, being translated at high efficiency. Indeed, counter to what was observed in genes with low TE, genes with high TE showed no significant change in TE following Tor-OE (Fig. 4H). Together, this analysis reveals that translation differs in dramatic ways from overall transcriptional expression, reflecting a highly dynamic translational landscape in the muscle.

**Fig 4:**
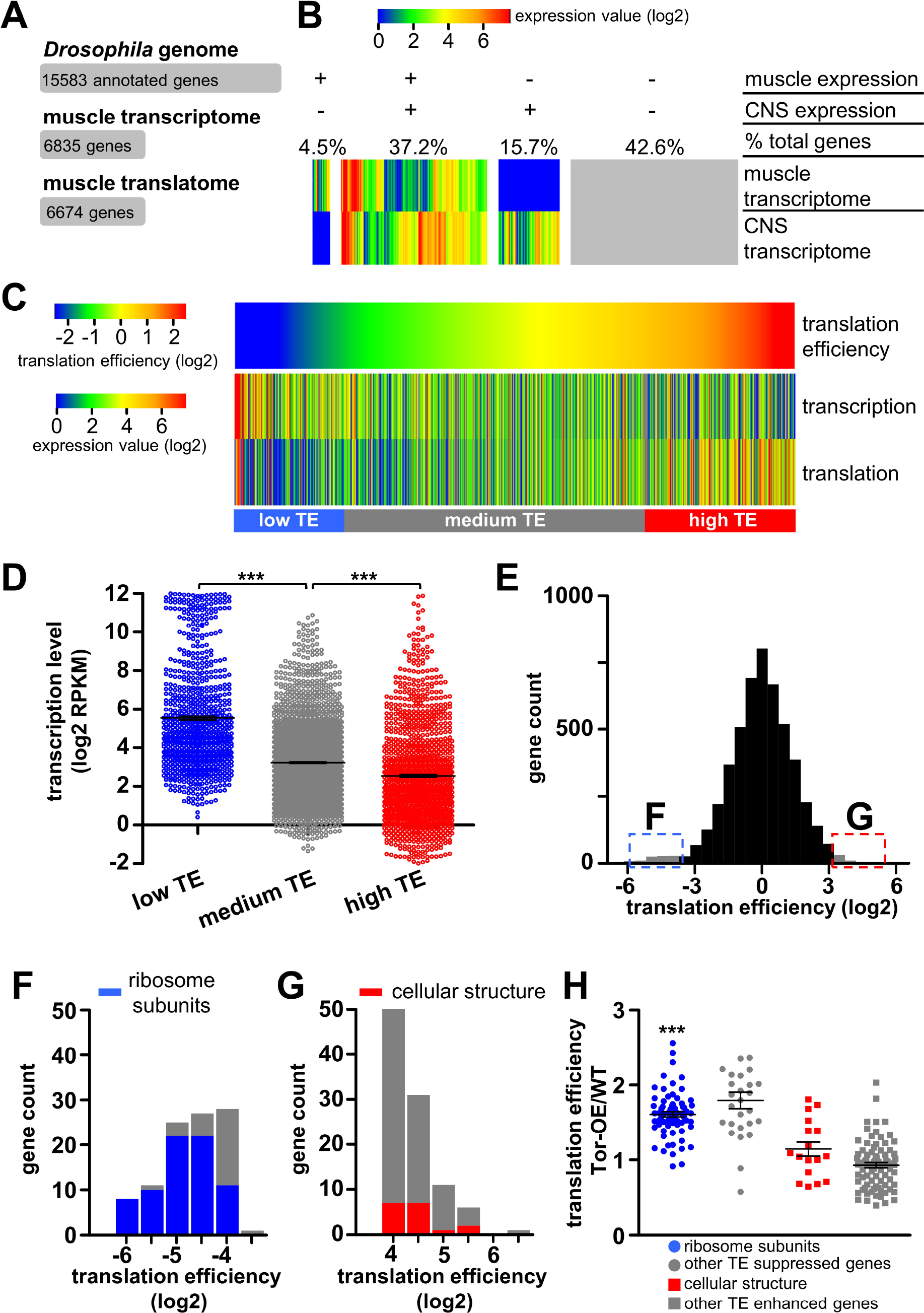
Analysis of the transcriptome and translatome reveals dynamic translational regulation in *Drosophila* muscle. **(A)** Definition of number of genes encoded in the *Drosophila* genome and those expressed in the muscle transcriptome and translatome. **(B)** Heatmap showing transcriptional levels of all annotated genes in the *Drosophila* larval muscle compared to those expressed in the central nervous system (CNS; (Brown et al., 2014)). These genes are grouped into four sections according to their expression status in muscle and CNS; the percentage of total genes is indicated above each section. **(C)** Heatmap showing translation efficiency (TE) and transcription and translation expression levels (RPKM) of genes expressed in muscle. Genes are ordered according to translation efficiency, with a trend observed for genes with high translation efficiency having low transcriptional expression levels and vice versa. **(D)** Transcriptional expression levels of genes with low TE (TE<0.5, blue), medium TE (TE between 0.5 and 2, grey) and high TE (TE>2, red). The transcriptional expression levels of genes in the low TE group is significantly higher than that of the medium TE group, while transcriptional expression of the high TE group is significantly lower than the that of the medium TE group (***=p<0.001; one-way ANOVA with post hoc Bonferroni’s test). **(E)** Histogram of translation efficiency across all genes expressed in the muscle. The 100 genes with the lowest translation efficiency (blue) and highest translation efficiency (red) are indicated. **(F)** Histogram of translation efficiency for the 100 genes with the lowest translation efficiency. An enrichment in ribosomal proteins, indicated in blue, is observed. **(G)** Histogram of translation efficiency for the 100 genes with the highest translation efficiency. Genes in the most abundant functional class, encoding proteins involved in cellular structure, are indicated in red. **(H)** Graph showing the 100 genes with the highest or lowest translation efficiency, their TE in Tor-OE as a ratio of wild type. Note that the translational efficiency of ribosomal proteins in Tor-OE are significantly increased compared to wild type. ***=p<0.001; paired Student’s t-test. Additional details can be found in Table S3, Table S4, and Figure S2.

### Transcriptional and ribosome profiling confirms expected changes in *GluRIIA* mutants and Tor-OE

We next confirmed the fidelity of our transcriptional and ribosome profiling approach by examining in molecular genetic detail the two manipulations we utilized to trigger postsynaptic retrograde signaling. The *GluRIIA^SP16^* mutation harbors a 9 kb deletion that removes the first half of the *GluRIIA* locus as well as the adjacent gene, *oscillin* (Fig. 5A; (Petersen et al., 1997)). Analysis of both transcriptional and ribosome profiling of *GluRIIA^SP16^* mutants revealed no transcription or translation of the deleted region, as expected (Fig. 5B). Transcription and translation of the adjacent gene, *oscillin*, was also negligible (wild type vs. *GluRIIA*: transcription=15.9 vs. 0.08 RPKM; translation=9.8 vs. 0.4 RPKM). However, the 3’ portion of the *GluRIIA* coding region was still transcriptionally expressed in *GluRIIA* mutants, while a significant reduction in translation was observed by ribosome profiling (Fig. 5B). Together, this confirms that although the residual 3’ region of the *GluRIIA* locus was transcribed, likely through an adjacent promoter, this transcript was not efficiently translated. Indeed, the peak ribosome profiling signals, which represent ribosome pause sites on the mRNA transcript, is known to be conserved for specific open reading frames (Li et al., 2012). However, this pattern was altered in *GluRIIA* mutants compared to wild type (Fig. 5B), suggesting the translation of the residual 3’ region of *GluRIIA* in *GluRIIA^SP16^* mutants was not in the same reading frame as the intact transcript. Thus, both transcriptional and ribosome profiling confirms that *GluRIIA* expression is abolished in *GluRIIA^SP16^* mutants.

**Fig 5:**
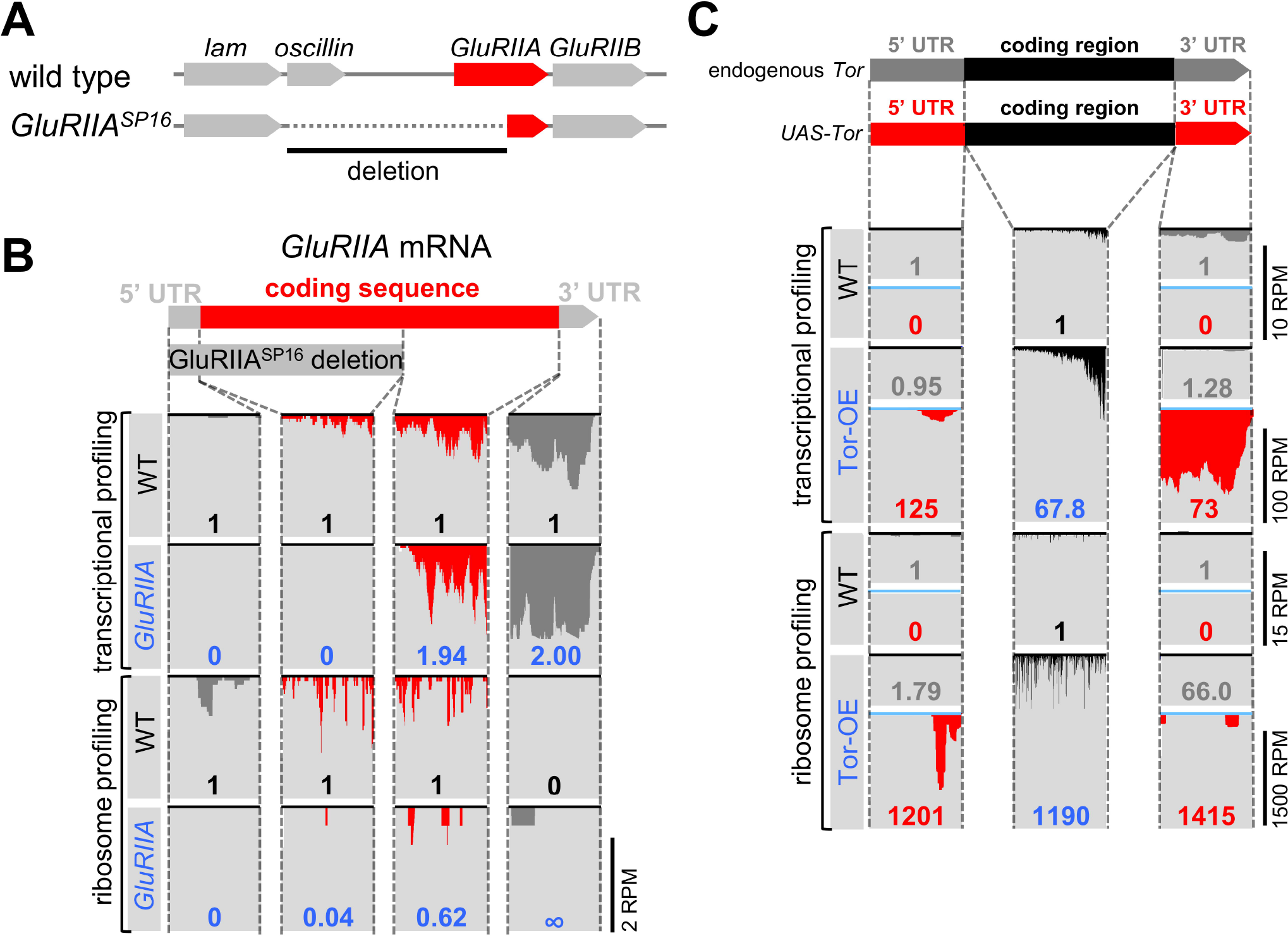
Transcriptional and translational profiling reveals expected changes in *GluRIIA* mutants and Tor-OE. **(A)** Schematic illustrating the genomic *GluRIIA* locus in wild type and *GluRIIA^SP16^* mutants. Note that the 5’ region of *GluRIIA* is deleted in the *GluRIIA* mutant, as well as the adjacent oscillin gene. **(B)** RNA-seq reads mapping to the *GluRIIA* locus from transcriptional and ribosome profiling in wild type and *GluRIIA^sp16^* mutants. The coverage graphs were divided into four sections corresponding to the regions indicated in the *GluRIIA* transcript. The numbers in each graph indicates the expression value of that region normalized to wild type transcriptional or ribosome profiling expression value. Note that no expression was detected by transcriptional or ribosome profiling in the deleted region in *GluRIIA* mutants, as expected. **(C)** Schematic illustrating the endogenous *Tor* mRNA transcript and the mRNA transcript encoded by Tor-OE. Both transcripts share the same coding sequence, but differ in their 5’UTR and 3’UTR sequences. Below are reads mapping to the indicated regions, divided into the three indicated sections. Note that both transcriptional and translational expression of *UAS-Tor* mRNA are significantly increased in Tor-OE, while transcription and translation of endogenous Tor mRNA are largely unchanged in Tor-OE.

Finally, we examined the expression of endogenous (genomic) and transgenically overexpressed (UAS) *Tor* through transcriptional and ribosome profiling. While both endogenous *Tor* and *UAS-Tor* mRNA share the same coding region, the 5’UTR and 3’UTR regions differ between these transcripts (Fig. 5C), enabling us to distinguish expression between these transcripts. We first confirmed a large increase in *Tor* coding region expression through both transcriptional profiling (68-fold) and ribosome profiling (1200 fold) (Fig. 5C, black). In contrast, analysis of the 5’ and 3’ UTRs revealed very little difference in endogenous *Tor* expression in *UAS-Tor* compared to wild type (Fig. 5C, grey), while a dramatic increase in both transcription (125-fold) and translation (1200-fold) was observed (Fig. 5C, red). Indeed, the translation efficiency of *Tor* was increased 14 fold in Tor-OE, consistent with the known influences of engineered 5’UTR and 3’UTR sequences in promoting translation in *UAS* constructs (Brand and Perrimon, 1993). Together, these experiments demonstrate that both transcriptional and ribosome profiling reliably report the expected changes in transcription and translation in the *Drosophila* larval muscle, and further serve to validate this approach.

### Transcriptional and ribosome profiling reveals post-translational mechanisms drive retrograde signaling

Given the substantial evidence that Tor-mediated control of new protein synthesis in the postsynaptic cell is necessary for retrograde PHP signaling (Penney et al., 2012), we compared transcriptional and translational changes in muscle between wild type, *GluRIIA* mutants, and Tor-OE. We anticipated a relatively small number of transcriptional changes, if any, between these genotypes, while we hypothesized substantial differences in translation rates would be observed both *GluRIIA* mutants and Tor-OE. The elevated translation of this exceptional subset of targets would, we anticipated, initiate postsynaptic PHP signaling and lead to an instructive signal that drives the retrograde enhancement in presynaptic release. Alternatively, we also considered the possibility that Tor-mediated protein synthesis may act in a non-specific manner, increasing overall protein synthesis in the postsynaptic cell, while there would be no overlap in translational changes between *GluRIIA* mutants and Tor-OE. In this case, post-translational mechanisms would operate on a global elevation in protein expression in Tor-OE, sculpting the proteome into an instructive retrograde signal. Indeed, the acute pharmacological induction and expression of PHP does not require new protein synthesis (Frank et al., 2006), providing some support for this model. We therefore compared transcription and translation in wild type, *GluRIIA* mutants, and Tor-OE.

We first compared transcription and translation in *GluRIIA* mutants and Tor-OE relative to wild type by plotting the measured expression values for each condition and determining the coefficient of determination, *r*^2^. An *r*^2^ value equal to 1 indicates no difference between the two conditions, while a value of 0 implies all genes are differentially expressed. This analysis revealed a high degree of similarity between wild type and *GluRIIA* mutants in both transcription and translation, with *r*^2^ values above 0.98 (Fig. 6A). In contrast, a slightly larger difference exists in transcription between Tor-OE and wild type, with *r*^2^=0.920 (Fig. 6B). Although transcription should not be directly affected by Tor-OE, this implies that perhaps an adaptation in transcription was induced in the muscle in response to chronically elevated translation. Finally, translational differences were the largest between Tor-OE and wild type, with *r*^2^=0.363 (Fig. 6B). This global analysis demonstrates there are very few transcriptional and translational changes in *GluRIIA* compared to wild type, while moderate transcriptional and robust translational changes exist in Tor-OE.

**Fig 6:**
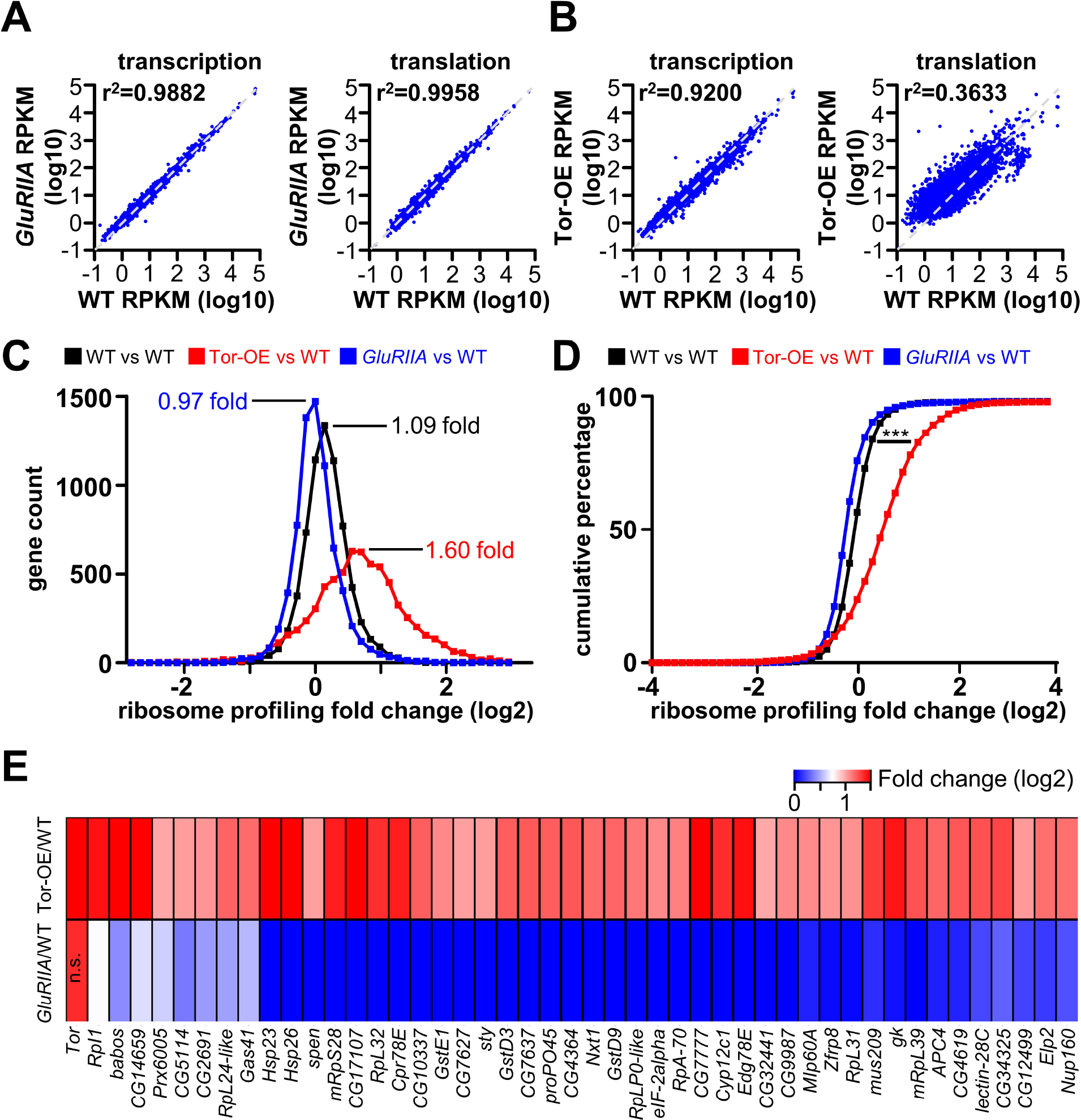
No changes in postsynaptic transcription or translation are observed in *GluRIIA* mutants. **(A)** Plot of transcriptional and translational levels of all expressed genes in *GluRIIA* mutants compared to wild type, with near identical correlations observed (indicated by *r*^2^ values). **(B)** Plot of transcriptional and translational levels of all expressed genes in Tor-OE compared to wild type. Note that while moderate changes in transcription are observed, large differences in translation are found (indicated by *r*^2^ values). **(C)** Histogram of the distribution of gene translation fold change in wild type versus wild type (black), which represents intrinsic variability, and that of Tor-OE versus wild type (Red), and that of *GluRIIA* mutants versus wild type (blue). Note the shift in distribution observed in Tor-OE, suggesting a global increase in translation. **(D)** Cumulative percentage plot of distributions shown in (C), showing significant difference between Tor-OE versus wild type distribution compared to wild type versus wild type distribution. (p<0.001, Kolmogorov–Smirnov test). **(E)** Heat map showing the 46 genes with significant increase in translation efficiency in Tor-OE, with the corresponding genes in *GluRIIA* mutants shown below. Note that no trend is observed in translational expression changes in these genes in *GluRIIA* mutants. Additional details can be found in Table S5.

In depth analysis of the transcriptome and translatome in *GluRIIA* muscle revealed that no genes were significantly altered. In particular, we eliminated genes that were up- or down-regulated due to known or expected influences in the genetic background (*GluRIIA* and *oscillin* expression, and closely linked genes to this locus; see Materials and Methods). Even with a lowered threshold for significant expression changes (fold change more than 2.5, with adjusted p-value less than 0.05), there were surprisingly no significant differences in either transcription or translation between *GluRIIA* and wild type. Given that Tor-OE both induced retrograde PHP signaling and drove a large and non-specific increase in translation (Fig. 6B; (Thoreen et al., 2012)), we considered whether a global shift in overall translation, and not specific translational regulation of particular targets, occurred in *GluRIIA* mutants, similar to what was observed in Tor-OE. First, we confirmed a global shift in translation in Tor-OE compared to wild type, as expected given the role of Tor as a general regulator of Cap-dependent translation initiation (Saxton and Sabatini, 2017). We plotted a gene count histogram of Tor-OE versus wild type measured by ribosome profiling, and overlaid the graph over a wild type over wild type ribosome profiling histogram. This analysis should indicate the relative degree of variation in translation between these groups. Indeed, a shift in global translation was observed in Tor-OE, with an average of 1.6 fold change in translation compared to 1.09 for wild type (Fig. 6C). This shift is significant when tested by Kolmogorov–Smirnov test (p<0.001) (Fig. 6D). We then performed this same analysis for *GluRIIA* vs WT. However, we observed no significant shift in translation in *GluRIIA* (0.97 fold change compared to 1.09; Fig. 6C). Thus, while Tor-OE induces a global increase in translation, loss of the *GluRIIA* receptor subunit in muscle does not measurably change overall translation.

Although no specific translational targets were identified to significantly change in *GluRIIA* mutants compared to wild type, we did identify 46 genes that exhibited significant increases in translation efficiency in Tor-OE (fold change more than 2, p-value less than 0.05) (Table S5). We characterized the expression of these genes in *GluRIIA* vs WT to determine whether a trend was observed that may differentiate the translational adaptations that drive retrograde PHP signaling in Tor-OE compared to the more general overall increase in protein synthesis. We therefore generated a heatmap of these 46 genes in Tor-OE vs WT and compared this to the same 46 genes in *GluRIIA* vs WT (Fig. 6E). This analysis revealed no particular trend or correlation in *GluRIIA* among the 46 genes with increased translation efficiency in Tor-OE (Fig. 6E). Together, these results suggest that retrograde signaling in the postsynaptic muscle, induced through loss of *GluRIIA,* does not alter translation of a specific subset of targets. In contrast, Tor-OE appears to induce a global increase in translation with no apparent specificity. This analysis indicates that while a similar retrograde enhancement in presynaptic release is induced by both loss of *GluRIIA* and Tor-OE, there is no overlap in translational targets, implying that post-translational mechanisms are ultimately required for retrograde homeostatic signaling.

### Chronic elevation in muscle protein synthesis leads to adaptive cellular responses

Although Tor-OE should only exert direct impacts on cellular translation, our analysis above indicated that transcriptional changes are induced following the global increase in translation by Tor-OE (Fig. 6B). This suggested that adaptations in transcription, and perhaps also translation, may have been triggered in Tor-OE in response to the cellular stress imparted by the chronic, global increase in muscle protein synthesis. Indeed, proteome homeostasis (proteostasis) is under exquisite control (Kong and Lasko, 2012; Vogel and Marcotte, 2012), and sustained perturbations in Tor activity induces transcriptional programs that adaptively compensate to maintain proteostasis (Tiebe et al., 2015; Wullschleger et al., 2006; Zhang et al., 2014). We therefore reasoned that by examining the changes in transcription and translation induced by Tor-OE, we may gain insight into how a cell adapts to the stress of chronically elevated translation.

Transcriptional and ribosome profiling revealed 11 genes with significantly upregulated transcription (fold change>3 and adjusted p-value<0.05; Fig. 7A and Table S6), and 75 genes with significantly upregulated translation (fold change>3 and adjusted p-value<0.05; Fig. 7A and Table S6) in Tor-OE compared to wild type. Interestingly, 8 of these genes exhibited shared increases in both transcription and translation (Fig. 7A), with their translational fold change (revealed by ribosome profiling) being larger than would be expected by their transcriptional fold change alone. This suggests a coordinated cellular signaling system that adaptively modulates both transcription and translation in response to the global elevation in translation following overexpression of Tor in the muscle. Further analysis revealed these upregulated genes to belong to diverse functional classes (Fig. 7B). Notably, we observed a striking enrichment in heat shock proteins and chaperones, factors known to assist with protein folding and participate in the unfolded protein response, particularly during cellular stress (Bukau et al., 2006; Hetz, 2012; Hohfeld et al., 2001; Romisch, 2005; Taipale et al., 2010). Indeed, among the 7 heat shock protein genes with significant expression in the muscle (Table S6), 5 were significantly upregulated in translation and 3 were significantly upregulated in transcription, with the remaining 2 showing a strong trend towards upregulation (Fig. 7C,D and Table S6). Given the well documented role for heat shock proteins in regulating protein folding, stability, and degradation in conjunction with the proteasome system (Hohfeld et al., 2001; Romisch, 2005; Taipale et al., 2010), this adaptation likely contributes to the stabilization of elevated cellular protein levels resulting from Tor-OE. Thus, the coordinated upregulation of heat shock proteins is one major adaptive response in transcription and translation following Tor-OE.

**Fig 7:**
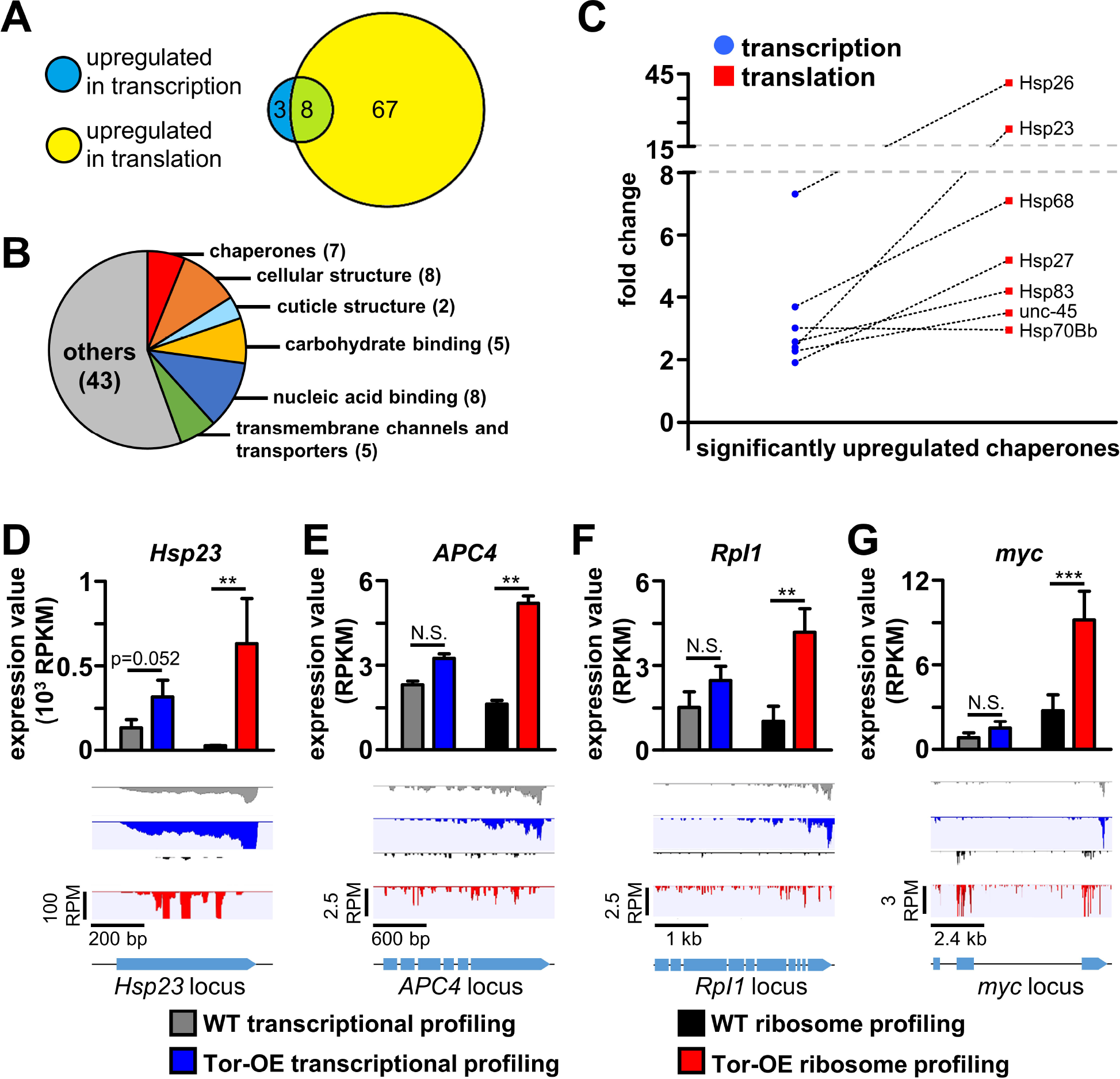
Increased cellular translation triggers adaptive cellular responses in both transcription and translation. **(A)** Diagram showing the number of significantly upregulated genes in transcription and translation in Tor-OE compared to wild type. **(B)** Pie chart showing the number of differentially upregulated genes in Tor-OE compared to wild type. **(C)** Graph showing transcriptional and translational changes for chaperones significantly upregulated in Tor-OE compared to wild type. Note that all but one exhibit higher translational changes compared to transcriptional changes, implying an additional upregulation in translational efficiency when compared to the increased transcriptional expression. Graph showing RPKM values measured by transcriptional and ribosome profiling in wild type and Tor-OE for the representative heat shock protein *Hsp23* **(D)**, the ubiquitin E3 ligase *APC4* **(E)**, the RNA polymerase *Rpl1* **(F)**, and the transcription factor *myc* **(G)**. Read mapping of the indicated genes are shown below. **=p<0.01, ***=p<0.001; Student’s t-test. Additional details can be found in Table S6.

In addition to heat shock proteins, we also identified genes involved in other cellular functions that are upregulated in Tor-OE and appear to enable adaptive responses to elevated cellular protein synthesis. For example, the E3 ubiquitin ligase subunit *APC4,* involved in protein degradation (Glickman and Ciechanover, 2002; Huang and Bonni, 2016), was upregulated in Tor-OE (Fig. 7E). Interestingly, proteasome subunits were reported to be upregulated in cells with increased Tor activity (Zhang et al., 2014). We also identified the RNA polymerase subunit *rpl1* and transcription factor *myc* to be upregulated following Tor-OE (Fig. 7F,G). These genes promote ribosome biogenesis, with Rpl1 necessary to synthesize ribosomal RNA and Myc involved in promoting the expression of ribosome assembly factors (van Riggelen et al., 2010; White, 2005). Together, Rpl1 and Myc likely promote the generation of additional ribosomes to meet the increased demands of protein synthesis induced by Tor-OE, consistent with previous studies showing Tor inhibition leads to decreased RpI1 transcription (Mayer et al., 2004). Hence, transcriptional and ribosome profiling defined adaptations in gene expression and protein synthesis that maintain proteostasis following chronic elevation in protein synthesis.

## DISCUSSION

We have developed a tissue-specific ribosome profiling strategy in *Drosophila* and used this approach to reveal the transcriptional and translational landscapes in larval muscle. This revealed significant differences between overall transcriptional and translational expression, and illuminated specific classes of genes with suppressed or elevated translation rates relative to transcription. We went on to leverage this technology to define the transcriptional, translational, and post-translational influences in the postsynaptic muscle that drive the retrograde control of presynaptic efficacy. Unexpectedly, we found no evidence that specific changes in transcription or translation are necessary for retrograde signaling, indicating that post-translational mechanisms ultimately transform the loss of postsynaptic receptors and enhanced protein synthesis into instructive retrograde cues. Finally, we identified adaptive cellular responses, in both transcription and translation, to chronically elevated protein synthesis that promote protein stability. Together, this study demonstrates the power of ribosome profiling in *Drosophila,* and illuminates the complex interplay of transcription, translation, and post-translational mechanisms that adaptively modulate cellular proteome stability and trans-synaptic retrograde signaling.

### Ribosome profiling and translational regulation in *Drosophila*

We have developed a highly efficient affinity tagging strategy and optimized RNA processing to enable tissue-specific ribosome profiling in *Drosophila.* Ribosome profiling has major advantages over measuring total mRNA expression and ribosome-associated mRNA (translational profiling using TRAP). Profound differences can exist between transcriptional expression and actual protein synthesis of genes expressed in a tissue. RNA-seq of total mRNA (transcriptional profiling) does not capture translational dynamics (Liu et al., 2016; Mortazavi et al., 2008). Translational profiling using TRAP does provide insights into translation (Heiman et al., 2014), but is less sensitive in detecting translational dynamics compared to ribosome profiling, which accurately quantifies the number of ribosomes associated with mRNA transcripts (Fig. 3; (Ingolia et al., 2012)). One major obstacle that limited the development of tissue-specific ribosome profiling is the relatively large amount of starting material necessary to generate the library for next generation sequencing. Because only ~30 nucleotides of mRNA are protected from digestion (Ingolia et al., 2009), ribosome profiling requires much more input material compared to standard RNA-seq (Brar and Weissman, 2015). Thus, the purification efficiency of the ribosome affinity tagging strategy and subsequent processing steps are very important to enabling successful profiling of ribosome protected mRNA fragments in *Drosophila* tissues. We achieved this high purification efficiency by systematically testing and optimizing multiple ribosome subunits (*RpL3, RpL36, RpS12, RpS13*) and affinity tags (6×His, 1×Flag, 3×Flag), finally settling on the *RpL3-3xFlag* combination to enable the highest purification efficiency (Fig. 2B and data not shown). Collectively, this effort differentiates our strategy from previous approaches in *Drosophila* that achieved ribosome profiling but lacked tissue specificity (Dunn et al., 2013) or purified ribosome-associated RNA from specific tissues but lacked the ability to quantify ribosome association with mRNA transcripts (Huang et al., 2013; Thomas et al., 2012; Yang et al., 2005; Zhang et al., 2016).

This optimized ribosome profiling approach has illuminated genome-wide translational dynamics in *Drosophila* muscle tissue and demonstrated two opposing protein production strategies utilized in these cells: elevated transcriptional expression coupled with low translation efficiency, which was apparent for genes encoding ribosomal subunits (Fig. 4F), and low transcriptional expression coupled with high translation efficiency, which was observed for genes encoding proteins belonging to diverse functional classes (Fig. 4G). These complementary strategies are likely tailored towards different cellular needs, enabling modulatory control of nuclear gene expression and cytosolic protein synthesis (Chekulaeva and Landthaler, 2016; Kong and Lasko, 2012). Thus, transcriptional and ribosome profiling of muscle tissue has revealed that translational control of ribosomal protein synthesis may be a phenomenon tailored to the unique metabolic needs of this tissue.

### Transcriptional, translational, and post-translational mechanisms required for retrograde synaptic signaling

We have used transcriptional and translational profiling to determine the contributions of transcriptional, translational, and post-translational mechanisms in the postsynaptic signaling system that drives the retrograde enhancement of presynaptic efficacy. Strong evidence has suggested that protein synthesis is modulated during homeostatic signaling at the *Drosophila* NMJ, with genetic disruption of Tor-mediated protein synthesis blocking expression and activation of the *Tor* pathway triggering expression (Kauwe et al., 2016; Penney et al., 2012; Penney et al., 2016). However, no specific translational targets in the muscle have been identified. We had expected that translational profiling would discover targets with increased translation efficiency in the muscles of *GluRIIA* mutants and/or following postsynaptic Tor overexpression, genetic conditions in which presynaptic homeostatic plasticity is chronically activated. However, no specific changes in transcription or translation were observed in *GluRIIA* mutants, while almost all muscle genes increased in translation following Tor-OE (Fig. 6). This implies that post-translational mechanisms ultimately drive PHP signaling in *GluRIIA* mutants. Furthermore, an apparent global increase in translation of nearly every gene also appears sufficient to instruct enhanced presynaptic release, consistent with the translational regulators implicated in PHP (Tor, S6 Kinase, eIF4E, and LRRK2), being non-specific cap-dependent translational regulators (Jackson et al., 2010; Penney et al., 2012; Penney et al., 2016). We consider several possible explanations and implications of these findings.

There are three conditions that trigger homeostatic retrograde signaling in the postsynaptic muscle: Acute pharmacological blockade of GluRIIA-containing postsynaptic receptors (Frank et al., 2006), genetic mutations in *GluRIIA* (Petersen et al., 1997), and chronic overexpression of Tor (Penney et al., 2012). First, all three manipulations lead to a similar enhancement in presynaptic release, and no additional increase in release was observed in *GluRIIA* mutants combined with Tor-OE (Penney et al., 2012). This indicates that these three perturbations may utilize the same signaling system; however, there is also evidence that post-translational mechanisms are necessary for PHP signaling. Indeed, the acute pharmacological induction of PHP does not require new protein synthesis (Frank et al., 2006), implying that if these manipulations use shared signal transduction and/or ultimately converge on the same pathway, post-translational mechanisms are responsible. Furthermore, there is evidence for post-translational mechanisms necessary for the induction of PHP signaling in *GluRIIA* mutants, as changes in CamKII phosphorylation and activity have been observed (Haghighi et al., 2003; Newman et al., 2017). In addition, other post-translational mechanisms, such as protein degradation or ubiquitination, could contribute to homeostatic signaling in the muscle. However, if all three manipulations do indeed ultimately utilize the same retrograde signal transduction system, it is quite intriguing that somehow the global increase in translation observed in Tor-OE is sculpted, perhaps by shared post-translational mechanisms, into a specific retrograde signal that instructs enhanced presynaptic release.

Second, it is possible that pharmacological, genetic, or Tor-OE-mediated inductions of PHP signaling are all mechanistically distinct, in which case no common transcriptional, translational, or post-translational mechanisms would be expected. Indeed, forward genetic screening approaches to discover genes necessary for PHP expression have failed to identify any genes needed for PHP induction in the postsynaptic muscle (Dickman and Davis, 2009; Muller et al., 2011), suggesting possible redundancy in these signaling systems. Finally, it is possible that very small, local changes in translation are necessary to drive retrograde signaling in *GluRIIA* mutants and Tor-OE, in which case our ribosome profiling approach may have lacked sufficient resolution to detect these changes, as tagged ribosomes were purified from whole cell muscle lysates. Indeed, a recent report demonstrated synapse-specific PHP expression (Newman et al., 2017), although this would not explain the translation-independent mechanism underlying PHP induction following pharmacological blockade of postsynaptic receptors. Future studies utilizing genetic, electrophysiological, biochemical, and imaging approaches will be necessary to identify the specific post-translational mechanisms that drive PHP signaling, and to what extent shared or distinct mechanisms are common between pharmacologic, genetic, and Tor-OE mediated PHP signaling.

### Proteostasis and adaptive cellular responses to elevated protein synthesis

Cells possess a remarkable ability to homeostatically control protein expression and stability, a process called proteostasis (Kaushik and Cuervo, 2015). This requires a robust and highly orchestrated balance between gene transcription, mRNA translation, and protein degradation (Sala et al., 2017; Vogel and Marcotte, 2012), while disruption of this process contributes to aging and disease (Hipp et al., 2014; Labbadia and Morimoto, 2015). Further, proteostatic mechanisms are not only customized to the unique demands of specific cells and tissues, but are adjusted throughout developmental stages and even tuned over hours according to diurnal metabolic and feeding cycles (Atger et al., 2015; Khapre et al., 2014; Sinturel et al., 2017; Wullschleger et al., 2006). The homeostatic nature of proteostasis is highlighted by the adaptations triggered in response to perturbations that threaten stable cellular protein levels, such as starvation and inhibitions of protein degradation (Bush et al., 1997; Fleming et al., 2002; Shang et al., 2011). We have used transcriptional and ribosome profiling to reveal new homeostatic adaptations triggered by proteostatic mechanisms that stabilize the proteome following chronic elevations in protein synthesis. In particular, genes that promote protein stability (chaperones), protein degradation, and ribosome biogenesis were transcriptionally and/or translationally upregulated following *Tor* overexpression in muscle, modulations in complementary pathways that synergistically prevent inappropriate protein interactions, promote protein removal, and increase the machinery necessary to maintain elevated protein synthesis (Claypool et al., 2004; Mayer et al., 2004; Zhang et al., 2014). Interestingly, many of these pathways are also targeted following other homeostatic perturbations to proteome stability, including heat shock, starvation, and inhibitions in protein degradation (Bar-Peled and Sabatini, 2014; Bush et al., 1997; Richter et al., 2010). This may suggest that proteostatic signaling involves a core program orchestrating adaptive modulations to transcription and translation to a diverse set of challenges to protein stability. Ribosome profiling enabled the definition of transcriptional and translational mechanisms that respond to chronic elevations of protein synthesis, revealing changes in translation that would not be apparent through profiling of total RNA expression alone.

Recent developments in next-generation sequencing have greatly expanded our ability to investigate complex biological phenomena on genome-wide scales. The power and variety of sophisticated genetic approaches are well-known in Drosophila. These include tissue-specific expression with a broad array of Gal4 and LexA drivers, transposable element manipulations, CRISPR/Cas-9 gene editing, and extensive collections of genetic mutations and RNAi lines (Gratz et al., 2015; Nagarkar-Jaiswal et al., 2015; Spradling et al., 1999; Venken and Bellen, 2014). Although some approaches have emerged to quantify RNA from entire tissues (Brown et al., 2014; Daines et al., 2011; White et al., 1999), as well as ribosome-associated RNA from specific tissues (Heiman et al., 2014; Huang et al., 2013; Sanz et al., 2009; Zhang et al., 2016), the technology described here now adds ribosome profiling to join this powerful toolkit to enable the determination of translation rates in specific cells at unprecedented resolution.

## MATERIALS AND METHODS

### Fly stocks and molecular biology

*Drosophila* stocks were raised at 25°C on standard molasses food. The *w^1118^* strain is used as the wild type control unless otherwise noted, as this is the genetic background of the transgenic lines and other genotypes used in this study. The following fly stocks were used: *GluRIIA^SP16^* (Petersen et al., 1997), *UAS-Tor-myc* (Wang et al., 2012), *RpL3^G13893^* (Bloomington Drosophila Stock Center, BDSC, Bloomington, IN, USA), *RpL3^KG05440^* (BDSC). All other *Drosophila* stocks were obtained from the Bloomington Drosophila Stock Center. Standard second and third chromosome balancers and genetic strategies were used for all crosses and for maintaining mutant lines. To control for the effects of genetic background on next generation sequencing data, we generated an isogenic stock. We then bred the genetic elements used in this study, (*BG57-Gal4, UAS-RpL3-Flag, GluRIIA^SP16^,* and *UAS-Tor-myc*) into this isogenic line and outcrossed for five generations to minimize the differences in the genetic background.

To generate the *UAS-RpL3-Flag* and *UAS-RpS13-Flag* transgenic lines, we obtained cDNA containing the entire coding sequences of *RpL3* (FBcl0179489) and *RpS13* (FBcl0171161). *RpL3* and *RpS13* coding sequence were PCR amplified and sub-cloned into the pACU2 vector (Han et al., 2011) with C-terminal 3xflag tag using a standard T4 DNA ligase based cloning strategy. To generate the genomic RpL3-3xflag construct, a 6.5kb sequence containing the entire *RpL3* genomic locus was PCR amplified from a genomic DNA preparation of *w^1118^* using the following primers 5’-ATCGGTACCACTTACTCCCTTGTTG-3’ and 5’-CAGCTGCAGGGTTTGTGACTGATCTAAAAG-3’. The same linker-3xflag sequence used in *UAS-RpL3-3xflag* was inserted right before the stop codon of *RpL3* of the PCR amplified genomic region using extension PCR. This sequence was cloned into the pattB vector (Groth et al., 2004). Constructs were sequence verified and sent to BestGene Inc. (Chino Hills, CA) for transgenic integration.

### Affinity purification of ribosomes and library generation

#### Tissue collection, lysis and library generation for transcriptional profiling

18 female third instar larvae were dissected in HL-3 saline as previously described (Chen et al., 2017), with all internal organs and the central nervous system removed, leaving only the body wall and attached muscle tissue. Following dissection, the tissue was immediately frozen on dry ice. The frozen tissue was then homogenized in 540 µl lysis solution (10 mM HEPES, PH 7.4, 150 mM KCl, 5 mM MgCl**2**, 100 µg/ml Cycloheximide) supplemented with 0.5% Triton-X100, 1U/µl ANTI-RNase (ThermoFisher scientific, AM2690) and protease inhibitor (EDTA-free, Sigma, COEDTAF-RO). 120 µl of the lysate was used for total RNA extraction by TRIzol LS Reagent (ThermoFisher scientific, 10296010). 2.5 µg of total RNA was used for isolation of mRNA with the Dynabeads mRNA DIRECT Purification Kit (ThermoFisher scientific, 61011). The entire isolated mRNA sample was used for library generation with the NEBNext Ultra Directional RNA Library Prep Kit for Illumina sequencing (NEB, E7420S).

#### Purification of ribosome associated mRNA and library generation (transcriptional profiling TRAP)

180 µl of the lysate described above was incubated with anti-flag antibody coated magnetic beads to purify ribosomes with associated mRNA. 75 µl of Dynabeads protein G (ThermoFisher scientific, 10004D) was used to coat 3 µg anti-Flag antibody (Sigma, F1804). The lysate-beads mixture was incubated at 4°C with rotation for 2 hours, then washed in buffer (10 mM HEPES, PH 7.4, 150 mM KCl, 5 mM MgCh_2_, 100 µg/ml Cycloheximide), supplemented with 0.1% Triton-X100 (0.1 U/µl SUPERase in RNase Inhibitor (ThermoFisher scientific, AM2696). RNA was extracted from ribosomes bound to the beads by TRIzol Reagent, and the co-precipitant linear acrylamide (ThermoFisher scientific, AM9520) was used to increase the RNA recovery rate. mRNA isolation and library generation were performed as described above.

#### Library generation for ribosome profiling

240 µl of lysate was incubated with anti-Flag antibody coated magnetic beads and 10000 units of RNase T1 (ThermoFisher scientific, EN0541) to perform digestion of exposed mRNA and ribosome purification simultaneously. 100 µl of Dynabeads protein G coated with 4 µg anti-Flag antibody was used. The lysate-beads-RNase T1 mixture was incubated at 4°C for 6 hours and washed; RNA was extracted as described above.

To perform size selection, the extracted RNA sample was separated on a denaturing 15% polyacrylamide urea gel. The gel region corresponding to 30-45 nt, as estimated by oligo markers, was excised. The gel slice was homogenized in 500 µl elution buffer (10 mM Tris-HCl, PH 7.5, 250 mM NaCl, 1 mM EDTA) supplemented with 0.2% SDS and RNAsecure reagent (ThermoFisher scientific, AM7005). The gel slurry was heated at 60°C for 10 min to allow inactivation of contaminating RNase by *RNAsecure* reagent and transferred to 4°C for overnight elution of RNA from the gel. The eluate was collected by centrifuging the gel slurry through a Spin-X column (Sigma, CLS8162), and RNA was precipitated by adding an equal volume of isopropanol and 25 µg linear acrylamide, incubated at room temperature for 30 min, and centrifuged for 15 min at 17000Xg, 4°C. The pellet was air dried and dissolved in 15 μl RNase-free water.

Library generation for ribosome profiling was performed using NEBNext Small RNA Library Prep Set for Illumina (NEB, E7330S) with minor modifications. The entire size selected mRNA fragments sample were first treated by phosphatase, rSAP (NEB, M0371S), to remove the 3’-phosphate. The samples were then incubated and denatured according to manufacturer’s instructions. RNA was precipitated from the reaction as described above, and the 3’ adaptor ligation was performed using NEBNext Small RNA Library Prep Set. The 5’-phosphate was then added to the mRNA fragments by supplying 2.5 μl 10 mM ATP, 1.5 µl 50 mM DTT and 0.5 µl T4 Polynucleotide Kinase (NEB, M0201S) to the 20 µl 3’ adaptor ligation reaction and incubating at 37°C for 30 min. 1 µl SR RT primer of the NEBNext Small RNA Library Prep Set was then added to the T4 polynucleotide kinase reaction and RT primer hybridization was performed. 5’ adaptor ligation, reverse transcription, PCR amplification and size selection of the PCR amplified library were performed using the NEBNext Small RNA Library Prep Set.

### High-throughput sequencing and data analysis

All libraries were sequenced on the Illumina NextSeq platform (single read, 75 cycles), and three replicates were performed for each genotype. Sequencing data analysis was performed using CLC genomics Workbench 8.0 software (Qiagen). Raw reads were trimmed based on quality scores, and adaptor sequences were removed from reads. Trimmed high quality reads were then mapped to the *Drosophila* genome (*Drosophila melanogaster,* NCBI genome release 5_48). Only genes with more than 10 reads uniquely mapped to their exons were considered reliably detected and further analyzed. Relative mRNA expression levels were quantified by calculating RPKM (Reads Per Kilobase of exon per Million mapped reads) using mapping results from transcriptional profiling. Relative translation levels were quantified by calculating RPKM (Reads Per Kilobase of exon per Million mapped reads) using mapping results from ribosome profiling. Translation efficiency was calculated by dividing ribosome profiling (or translational profiling TRAP) RPKM by transcriptional profiling RPKM.

To determine differentially transcribed or translated genes, a weighted t-type test (Baggerly et al., 2003) was performed based on three replicate expression values for each gene between *GluRIIA* mutants and wild type, and Tor-OE and wild type using the statistical analysis tool of CLC genomics workbench. The analysis was performed on expression values obtained by transcriptional profiling to determine differentially transcribed genes, and on expression values obtained by ribosome profiling to determine differentially translated genes. Genes with a p-value less than 0.05 and fold change higher than 3-fold were considered differentially transcribed or translated. We also determined differentially transcribed or translated genes using R package DESeq2 analysis (Love et al., 2014), considering genes with adjusted p-values less than 0.05 as differentially expressed. The Baggerly’s t test method and DESeq2 method produced highly similar lists of differentially expressed genes. Genes only revealed by one method were excluded from the final list. To determine gene targets undergoing translational regulation in *GluRIIA* mutants and Tor-OE compared to wild type, two criteria were used. First, the gene must have at least a 2-fold significant increase (p<0.05, Student’s t test) in translation efficiency compared to wild type. Second, a significant increase in ribosome profiling expression value (p<0.05, Baggerly’s t test) must also exist for the same gene. These two criteria ensure identification of genes that have true translational up-regulation that are not due to transcriptional changes.

### Immunocytochemistry and confocal imaging

Third-instar larvae were dissected in ice cold 0 Ca^2+^ HL-3 and fixed in Bouin’s fixative for 2 min as described (Chen et al., 2017). Mouse anti-Flag (Sigma, F1804) was used at 1:500, while donkey anti-mouse Alexa Fluor 488-conjugated secondary antibody (Jackson Immunoresearch) was used at 1:400. Alexa Fluor 647-conjugated goat anti-HRP (Jackson ImmunoResearch) was used at 1:200. Samples were imaged using a Nikon A1R Resonant Scanning Confocal microscope equipped with NIS Elements software and a 100x APO 1.4NA oil immersion objective, using separate channels with two laser lines (488 nm and 561 nm). Images were obtained using settings optimized for detection without saturation of the signal.

### Electrophysiology

All recordings were performed in modified HL-3 saline supplied with 0.3 mM Ca^2+^ as described (Chen et al., 2017; Kiragasi et al., 2017).

### Statistical Analysis

All data are presented as mean +/−SEM. Student’s *t* test was used to compare two groups. A one-way ANOVA followed by a post-hoc Bonferroni’s test was used to compare three or more groups. All data was analyzed using Graphpad Prism or Microsoft Excel software, with varying levels of significance assessed as p<0.05 (*), p<0.01 (**), p<0.001 (***), N.S.=not significant. Statistical analysis on next generation sequencing data was described in the High-throughput sequencing and data analysis section.

## ACKNOWLEDGEMENTS

The authors declare no competing financial interests. We thank Pejmun Haghighi (Buck Institute, CA, USA) for sharing *Drosophila* stocks. We thank Chun Han (Cornell University, NY, USA) for sharing the pACU2 vector and for insights into controlling gene expression with this vector. We acknowledge the Developmental Studies Hybridoma Bank (Iowa, USA) for antibodies used in this study, and the Bloomington Drosophila Stock Center (NIH P4OD018537) for fly stocks. This work was supported by a grant from the National Institutes of Health (NS019546) and research fellowships from the Alfred P. Sloan, Ellison Medical, Whitehall, Klingenstein-Simons, and Mallinckrodt Foundations to DKD.

## AUTHOR CONTRIBUTIONS

The project was conceived by XC and DKD. XC obtained all experimental data. The manuscript was written by XC and DKD.

---

**Fig S1:**
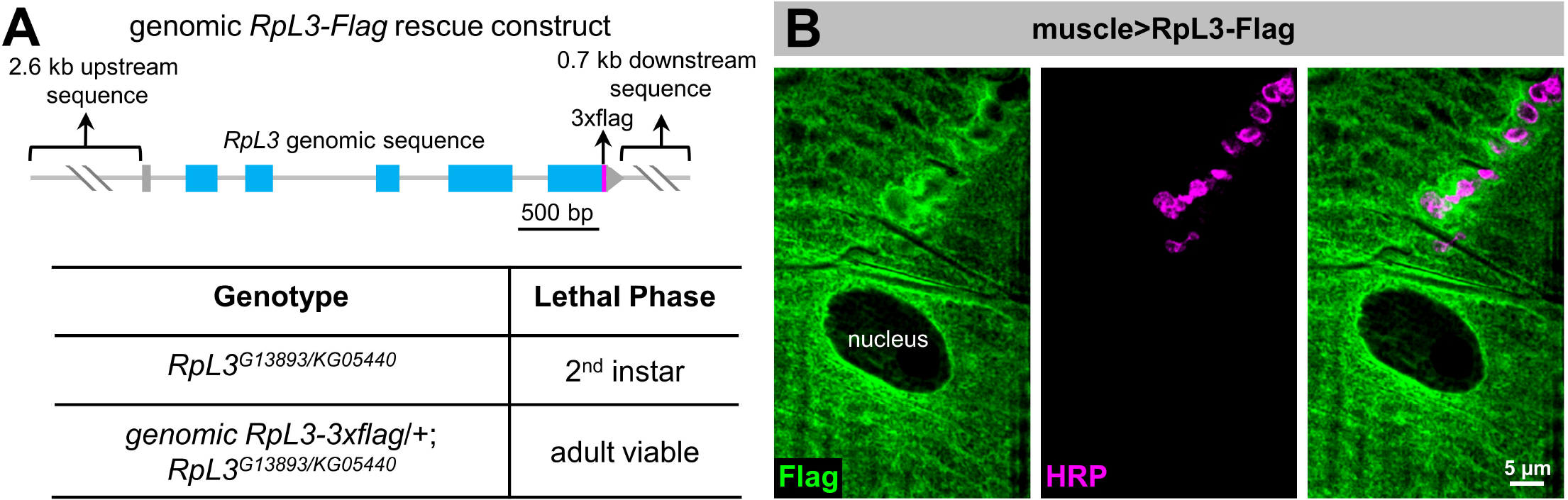
RpL3-Flag localization in muscle and rescue of *RpL3* mutants. **(A)** Schematic illustrating the genomic *RpL3-3xflag* rescue construct and table showing lethal phase of *RpL3* mutant (*w*;*RpL3*^*G13893/KG05440*^) and RpL3-Flag rescue (*w*;*genomic-RpL3-3xflag/+*;*RpL3*^*G13893/KG05440*^). **(B)** Representative images of RpL3-1216 Flag expressed in muscle Representative images of RpL3-1216 Flag expressed in muscle (*w*;*BG57-Gal4/UAS-RpL3-3xflag*) immunostained with anti-Flag (green) and anti-HRP (magenta) antibodies.

**Fig S2:**
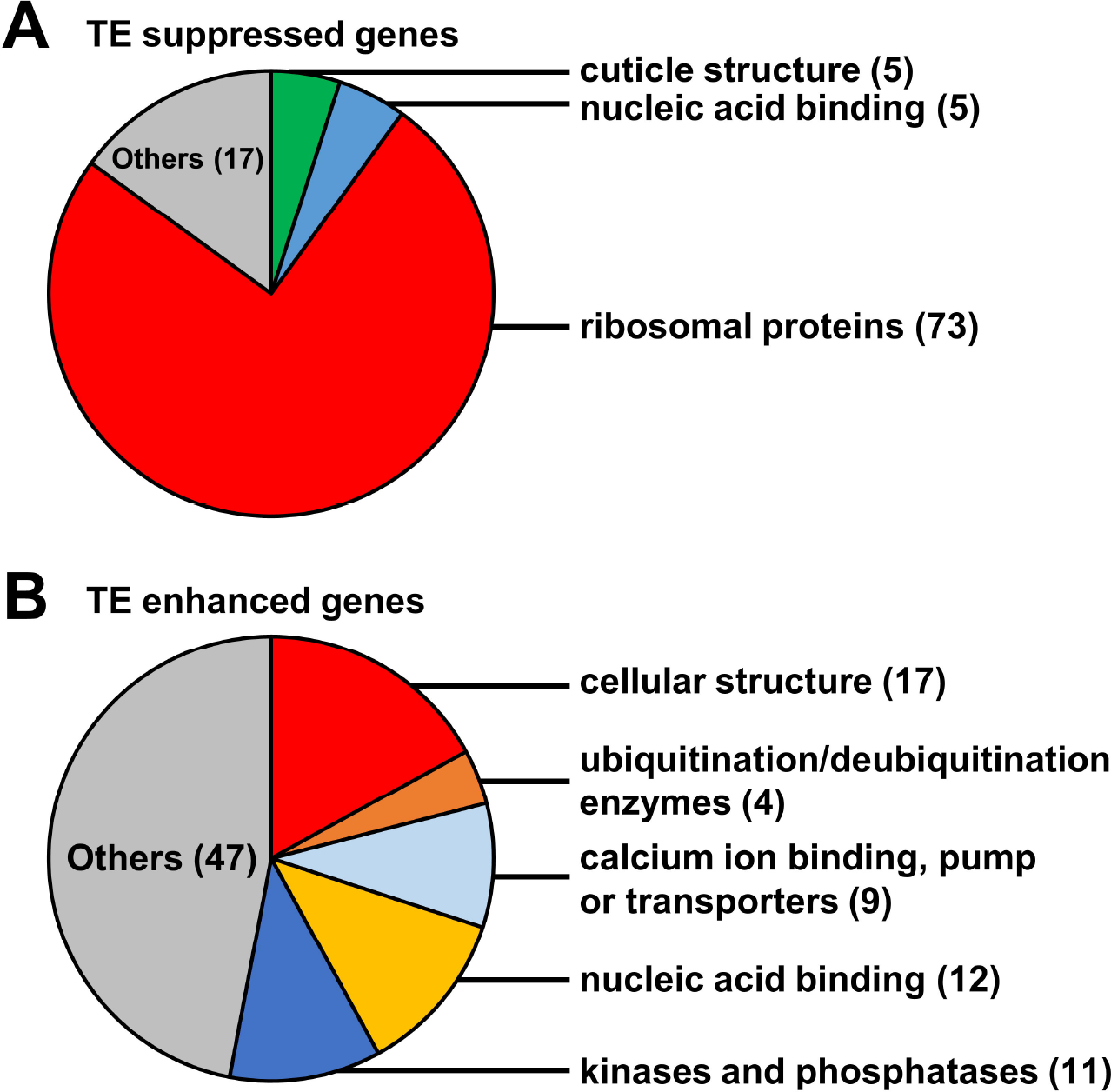
Functional classes for the 100 genes with the lowest and highest TE. **(A)** Functional classes for the 100 genes with the lowest TE. Note that ribosomal proteins represent the largest class, with 73 of the 100 genes encoding ribosomal proteins. **(B)** Functional classes for the 100 genes with the highest TE. Diverse functional classes are present, with genes encoding proteins involved in the cellular structure being the most abundant class.

**Table S1: Next generation sequencing data analysis statistics.**

**Table S2: Genes identified in the *Drosophila* third-instar larval transcriptome and translatome.** Genes in the transcriptome or translatome are defined by having at least 10 unique exon reads by transcriptional or ribosome profiling in all three replicates. Genes are listed in the order of transcription (transcriptional profiling RPKM). Genes only found in the transcriptome or translatome are listed at the end of the respective list.

**Table S3: Details on the 100 genes with the lowest translation efficiency.**

**Table S4: Details on the 100 genes with the highest translation efficiency.**

**Table S5: Details on the genes with significantly increased translation efficiency in Tor-OE vs wild type.**

**Table S6: Genes significantly up-regulated in transcription 1241 or translation in Tor-OE vs wild type.** Significantly up-regulated genes are defined by p-values<0.05 (see materials and methods) and fold changes>3.

